# DeepUMQA: Ultrafast Shape Recognition-based Protein Model Quality Assessment using Deep Learning

**DOI:** 10.1101/2021.10.02.462491

**Authors:** Sai-Sai Guo, Jun Liu, Xiao-Gen Zhou, Gui-Jun Zhang

## Abstract

**Motivation:** Protein model quality assessment is a key component of protein structure prediction. In recent research, the voxelization feature was used to characterize the local structural information of residues, but it may be insufficient for describing residue-level topological information. Design features that can further reflect residue-level topology when combined with deep learning methods are therefore crucial to improve the performance of model quality assessment.

**Results:** We developed a deep-learning method, DeepUMQA, based on Ultrafast Shape Recognition (USR) for the residue-level single-model quality assessment. In the framework of the deep residual neural network, the residue-level USR feature was introduced to describe the topological relationship between the residue and overall structure by calculating the first moment of a set of residue distance sets and then combined with 1D, 2D, and voxelization features to assess the quality of the model. Experimental results on test datasets of CASP13, CASP14, and CAMEO show that USR could complement the voxelization feature to comprehensively characterize residue structure information and significantly improve the model assessment accuracy. DeepUMQA outperformed the state-of-the-art single-model quality assessment methods, including ProQ2, ProQ3, ProQ3D, Ornate, VoroMQA, and DeepAccNet.

**Availability:** The source code and executable are freely available at https://github.com/iobio-zjut/DeepUMQA.

## 1 Introduction

Protein is the basis of life matter and ubiquitous in almost all biological processes. High-throughput acquisition of the protein structure helps in understanding its function and mechanism of action (Yang *et al*., 2015; Zhou *et al*., 2019a; Senior *et al*., 2020). Although the current sequencing technology can quickly and cheaply determine the primary sequence of a protein, determining the 3D structure of a protein is still difficult and expensive (Zhang *et al*., 2016; Zhou *et al*., 2019b). In the past few decades, many methods have been proposed to predict the 3D structure of a protein directly from the primary sequence (Rohl *et al*., 2004; Leaver-Fay *et al*., 2011; Xu *et al*., 2012; Moult *et al*., 2018; Liu *et al*., 2020; Zheng *et al*., 2021). In particular, the introduction of deep residual neural networks has promoted the rapid development of structural prediction (Wang *et al*., 2017; AlQuraishi *et al*., 2019; Kryshtafovych *et al*., 2019; Kuhlman *et al*., 2019; Xu *et al*., 2019; Mao *et al*., 2020; Yang *et al*., 2020). In CASP14, the predicted model of AlphaFold2 in most target proteins is comparable with that of experimental structures (Jumper *et al*., 2021). Assessing the quality of the generated protein model is a fundamental part of protein structure prediction, which is important for further structure refinement and reliable identification of the best models (Shuvo *et al*., 2020; Cheng *et al*., 2019).

In general, protein model assessment methods can be divided into two categories. The first covers single-model quality assessment approaches, which take a single structural model as an input, extract features that can reflect model information, and use machine learning methods to infer the quality of the model (Ray *et al*., 2012; Uziela *et al*., 2016; Olechnovic *et al*., 2017; Pagès *et al*., 2019; Jing *et al*., 2021; Hiranuma *et al*., 2021). The second one encompasses consensus methods, which evaluate protein models by using information from other models in a pool of candidate models (Lundström *et al*., 2001; Ginalski *et al*., 2003; Cheng *et al*., 2009). Although consensus methods achieve high correlations between predicted and true quality measures, their performance is greatly affected by the size and diversity of the input model pool (Cao *et al*., 2017). When the models lack consensus or are similar, the consensus method cannot easily select the best model (Shuvo *et al*., 2020). By contrast, single-model quality assessment methods are not limited by the model pool and can independently score and select models. Single-model quality assessment methods have elicited increasing attention in the recent Critical Assessment of Techniques for Protein Structure Prediction (CASP) (Manavalan *et al*., 2017; Kryshtafovych *et al*., 2018; Cheng *et al*., 2019; McGuffin *et al*., 2019). In CASP14, single-model quality assessment methods account for more than 70% of all model quality assessment methods (Kwon *et al*., 2021).

Single-model quality assessment methods usually use different feature combinations and machine learning methods to learn the implicit relationship between a model’s structure and its quality. ProQ2 predicts the local and global quality of protein models by using support vector machine (SVM) and weighting the sequence spectra of residue specific features including atom-atom contacts, residue-residue contacts, and surface area, predicted and observed secondary structures (Ray *et al*., 2012). ProQ3 uses Rosetta energy terms, which are based on the fullatom model and simplified centroid side chain representation (centroid model), as input features to train the same SVM employed in ProQ2 (Uziela *et al*., 2016). VoroMQA combines the idea of statistical potentials with the use of interatomic contact area, which is adopted to describe and seamlessly integrate explicit interactions between protein atoms and implicit interactions of protein atoms with the solvent (Olechnovic *et al*., 2017). With the introduction of the convolutional residual neural network (ResNet) into protein structure prediction, an increasing number of singlemodel quality assessment methods utilize deep learning (Xu *et al*., 2019). ProQ3D replaces SVM in ProQ3 with multilayer perceptron, which significantly improves the accuracy of model assessment (Uziela *et al*., 2017). Ornate predicts local (residue-wise) and global model quality based on 3D voxel atomic representation and 3D convolutional network. The input density map corresponding to each residue and its neighborhood is aligned with the backbone topology of this residue as the voxelization feature of residues that are translationally and rotationally invariant (Pagès *et al*., 2019). DeepAccNet uses 3D and 2D convolution to predict the per-residue accuracy and residue-residue distance signed error in protein models and employs these predictions to guide Rosetta protein structure refinement, in which 3D convolution similar to that in Ornate is used to evaluate the local atomic environment, and 2D convolution is adopted to provide the global context (Hiranuma *et al*., 2021). ProteinGCN represents the initial model as a graph, in which atoms, residues, and geometric features are first extracted; then, a graph neural network (GNN) is subsequently used to predict the quality of the protein model (Sanyal *et al*., 2020). GNNRefine applies GNN to predict the refined inter-atom distance probability distribution of the input model for further protein model refinement (Jing *et al*., 2021). Existing methods show that feature extraction and network model training are two key factors that affect the performance of model quality assessment. Designing input features that reflect structural information in multiple dimensions helps in inferring the mapping relationship between the structure of the model and quality.

In this work, we developed an Ultrafast Shape Recognition (USR)-based residue-level single model quality assessment method, termed DeepUMQA. The residue-level USR feature is used to map the topological information of the model to each residue, thereby characterizing the relationship between the residue and the topological structure (i.e., the topological information of the residue). Under the framework of the deep residual neural network, the USR feature is combined with the residue voxelization feature (i.e., the local structure information of the residue), secondary structure, distances, and Rosetta energy terms to predict the quality of the protein model. Experimental results show that the USR feature is complementary to the voxelization feature in describing residues from local and topological aspects, and it can thus significantly improve the performance of model quality assessment.

## 2 Methods

The pipeline of DeepUMQA is shown in Figure 1. It consists of four steps, namely, data preparation, feature extraction, network architecture, and model training and lDDT calculation for model quality assessment.

**Fig. 1.**
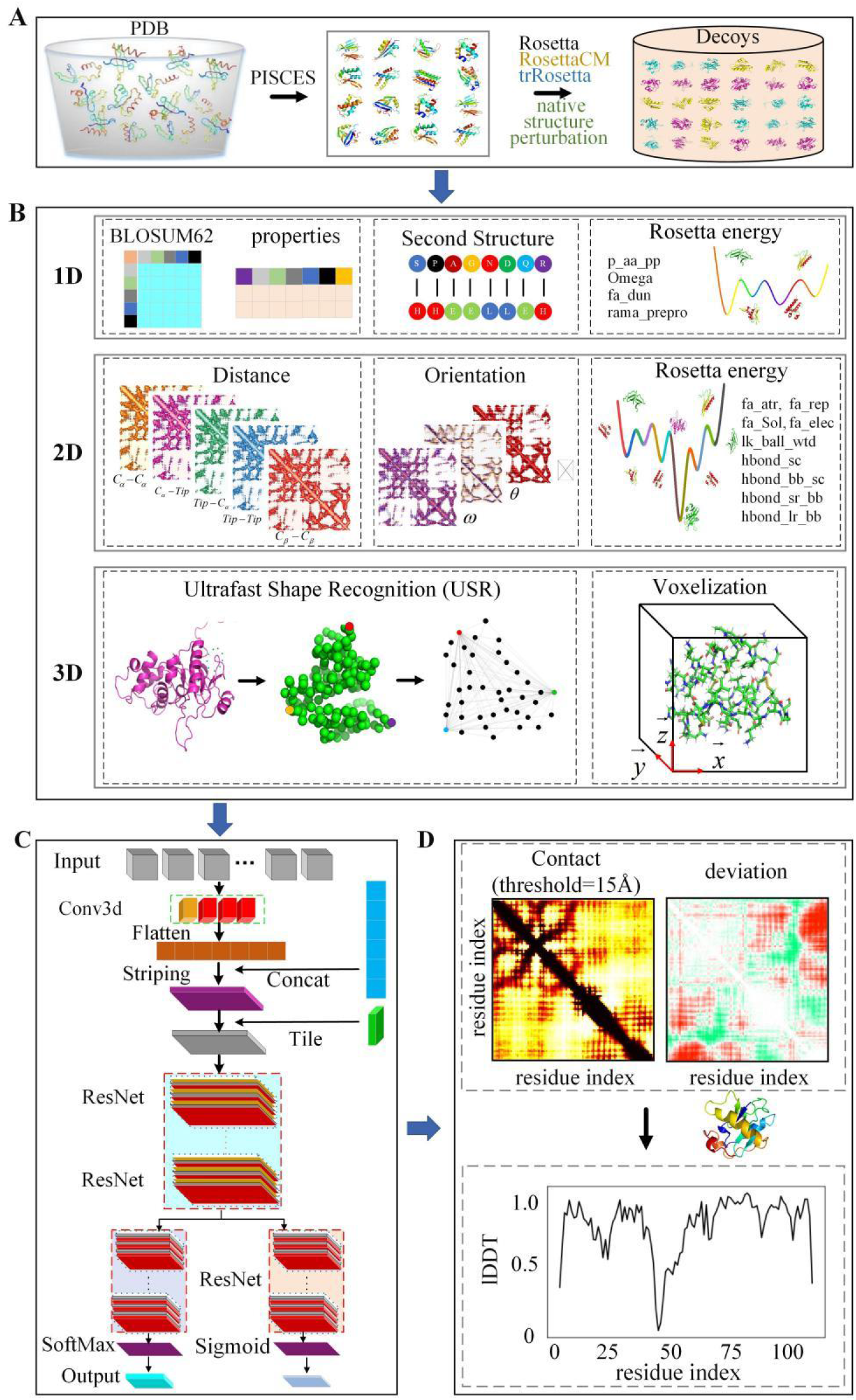
Pipeline of DeepUMQA. (A) Data preparation. (B) Feature extraction. (C) Network architecture. (D) Model training and lDDT calculation.

### 2.1 Data preparation

We construct a non-redundant protein dataset from the PISCES server (Wang *et al*., 2003), and the filter criteria are set as follows: (i) maximum sequence redundancy of 40%, (ii) minimum resolution of 2.5Å, (iii) each protein chain is limited to 50-300 residues, and (iv) the protein be either a monomer or interact with other chains with minimum (less than 1 kcal/mol of Rosetta energy). The dataset contains 7,615 proteins (7,226 for training and 389 for validation). Four methods including Rosetta stochastic modeling (Rohl *et al*., 2004), comparative modeling from RosettaCM (Song *et al*., 2013), native structure perturbation, and folding guided by the deep learning method trRosetta (Yang *et al*., 2020) are used to generate structure models with variety and a reasonable precision distribution. Each protein generates about 150 decoys, and each decoy is relaxed through Rosetta dual-relax (Conway *et al*., 2014). To test the performance of DeepUMQA, we collect a test dataset consisting of 51 CASP13, 44 CASP14, and 195 CAMEO targets. The detailed information of these targets are listed in Tables S5-S7 in Supplementary Information (SI). Each target of CASP13 and CASP14 obtains about 150 models from the official website (https://predictioncenter.org/download_area/), and each target of CAEMO obtains about 16 models from the official website (https://www.cameo3d.org/quality-estimation/), thereby forming a dataset containing 17,057 models in total.

### 2.2 Features extraction

As shown in Figure 1(B), 1D features (i.e., amino acid sequence properties, secondary structure, and Rosetta intra-residue energy terms), 2D features (i.e., residue-residue distance and orientation and Rosetta inter-residue energy terms), and 3D features (i.e., USR and voxelization at the residue level) are extracted. All features are listed in Table S1 in SI.

#### 2.2.1 Ultrafast Shape Recognition (USR)

Although the Cartesian coordinates of all atoms can completely describe the structural information of the protein model, this representation changes with rotation, which greatly increases the complexity of network training and usage (Hiranuma *et al*., 2021). To describe the local structure information of residues, previous methods, such as Ornate (Pagès *et al*., 2019) and DeepAccNet (Hiranuma *et al*., 2021), voxelize each residue individually in the corresponding local coordinate frame defined by the backbone *C, C*_*α*_, and *N* atoms. Such representation is translationally and rotationally invariant because projections onto local frames are independent of the global position of the protein structure in 3D space. The voxelization method effectively describes the local structure information of the residue, but it does not fully reflect the topological relationship between the residue and the overall structure. Moreover, the calculation of the voxelization feature vector and 3D convolution are extremely complicated and time consuming.

Inspired by the Ultrafast Shape Recognition method (Ballester *et al*., 2007; Hao *et al*., 2016), the topological information of the protein structure can be quickly captured with almost no additional computational cost by selecting an appropriate set of interatomic distances. In particular, this subset can be selected as the set of all atom distances from a reduced number of strategic reference locations. Figure 2 shows a schematic of using USR to characterize protein topological structure information. Four reference locations that effectively represent the center and boundary of the protein structure are considered, and a subset of distances between them is utilized to quickly identify the topology of the protein.

**Fig. 2.**
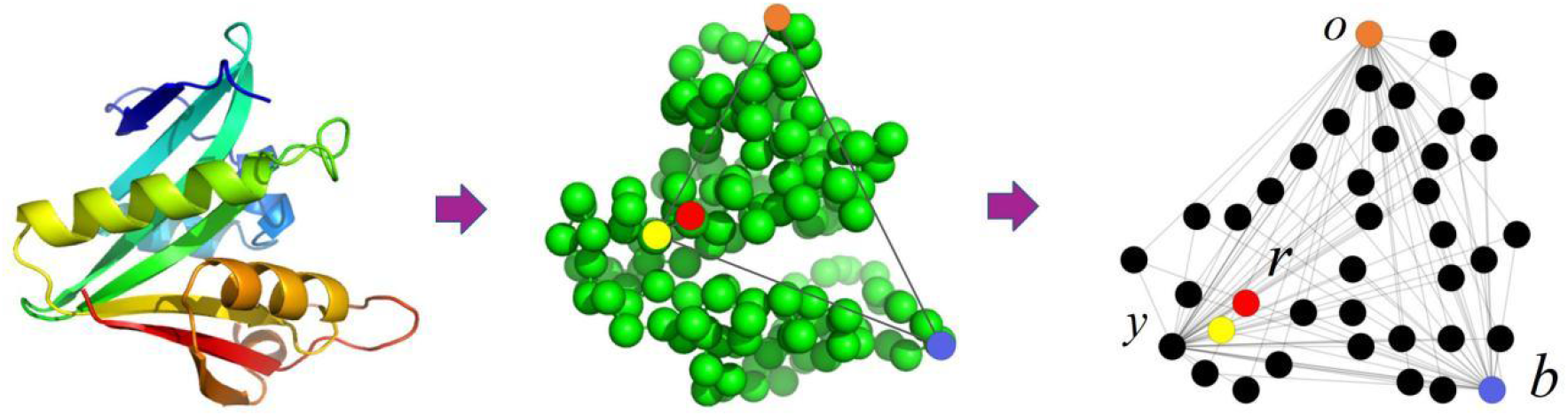
Schematic of the structures of proteins using USR. First, four reference atoms are considered: the protein structure centroid *r* (red dot), the atom *y* (yellow dot) closest to the centroid, the atom *b* (blue dot) farthest from the centroid, and the atom *o* (orange dot) farthest from atom *b*. Second, the distances of the four selected reference atoms to all other atoms in the structure are calculated to form a 4-dimensional distance set. Lastly, in accordance with the distance set, their first moments are calculated to describe the entire protein structure.

In this work, we introduce USR to the residue level to describe the topological relationship between the residues and the overall structure. For a given protein structure model, we extract the USR features of each residue. As shown in Figure 3, for the *i*th residue, with the current residue *r*_*i*_ as a reference, the residue *b*_*i*_ farthest from the residue *r*_*i*_ and the residue *g*_*i*_ farthest from the residue *b*_*i*_ are determined successively. The topological relationship between the current residue and the overall structure can be described by the three residue distance sets defined by *r*_*i*_, *b*_*i*_, *g*_*i*_. For residue *r*_*i*_, distances from it to all other residues in the structure are calculated to form a distance set. For residues bi and *g*_*i*_, the same method is applied to obtain the corresponding distance set. The distance between residues is calculated according to the distance between the *C*_*β*_ (*C*_*α*_ for glycine) atoms of the residues. The first moment 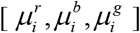 of the residue distance sets is calculated as the feature mapping of the current residue topology information, and 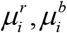, and 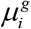 are defined as follows:

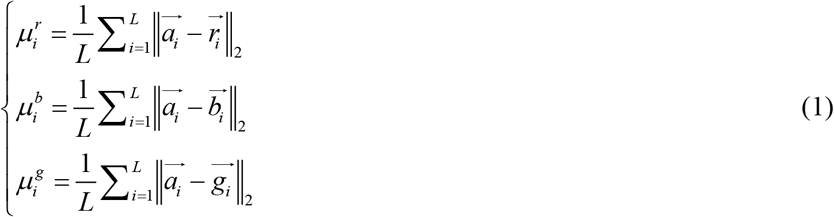

where *L* is the sequence length of the protein, 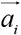 is the coordinate of the *C*_*β*_ (*C*_*α*_ for glycine) atom of the ith residue, and 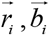, and 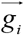 are the coordinates of the *C*_*β*_ (*C*_*α*_ for glycine) atoms of the residues *r*_*i*_, *b*_*i*_, and *g*_*i*_, respectively.

**Fig. 3.**
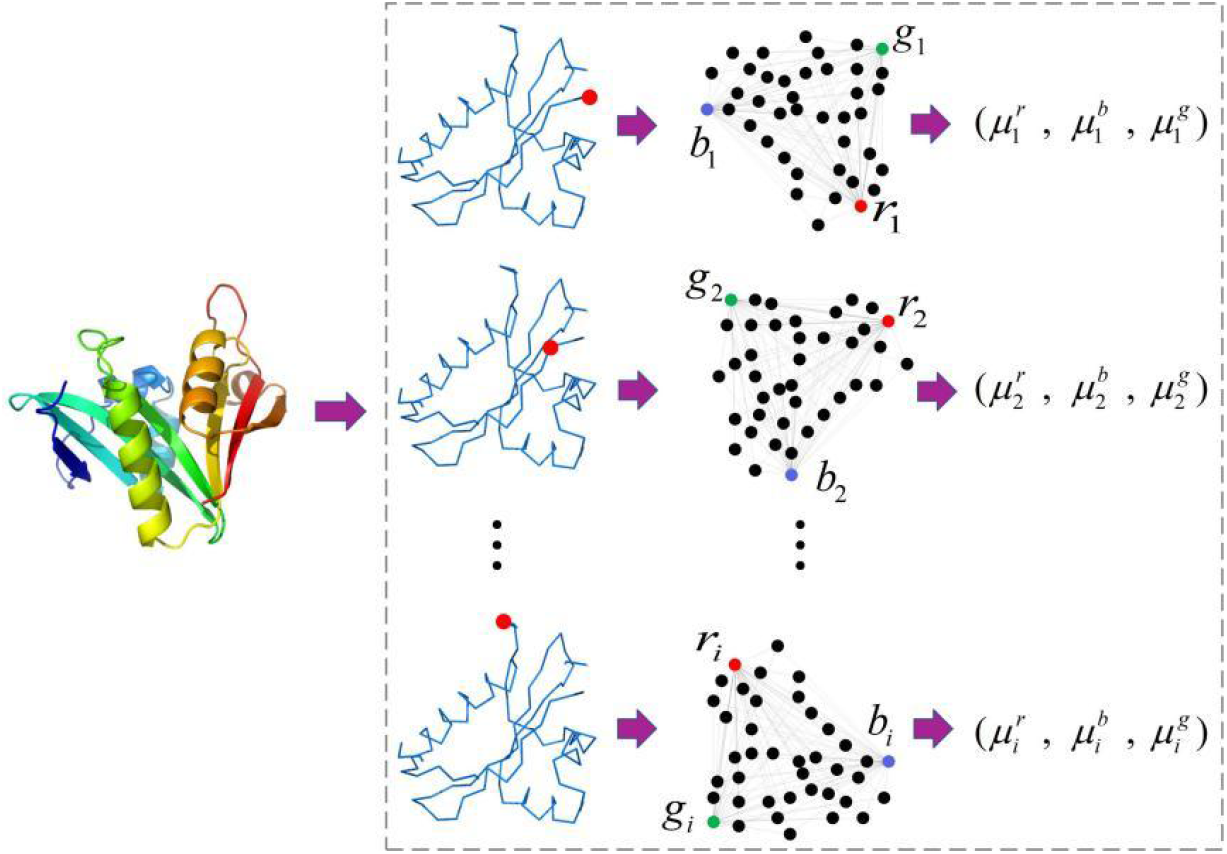
Residue level USR feature of protein structure. 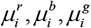 are the results of the first moment calculation of the distance sets at the three reference positions: the current residue *r*_*i*_ as a reference, the residue *b*_*i*_ farthest from the residue *r*_*i*_ and the residue *g*_*i*_ farthest from the residue *b*_*i*_.

#### 2.2.2 Amino acid sequence properties and secondary structure

The amino acid sequence contains the 3D structure information of the protein, thus, the properties of the amino acid sequence are important for inferring the quality of the structure model. Blosum62 is a kind of amino acid substitution scoring matrix commonly used in sequence comparison in bioinformatics at present (Henikoff *et al*., 1992). It is used to derive the replacement vector of the residue at each position of the structure, thus obtaining an input feature. In addition, we also obtain per amino-acid feature sets from Meiler as the features of individual residues (Meiler *et al*., 2001). The secondary structure information is also useful for describing the structure of proteins. The DSSP is utilized to obtain the secondary structure characteristic information of the subsequence of the protein structure (Kabsch *et al*., 1983). These feature items are quantified before being used. The per amino acid feature sets from Meiler and Blosum62 scoring matrix are listed in Tables S2 and S4 in SI, respectively.

#### 2.2.3 ROSETTA energy terms

In this work, 13 Rosetta centroid energy terms are also applied as features(Rohl *et al*., 2004; Leaver-Fay *et al*., 2011), including one-body-terms (omega, p_aa_pp, fa_dun, and rama_prepro), two-body-terms (fa_atr, fa_rep, fa_sol, lk_ball_wtd, fa_elec, hbond_bb_sc, and hbond_sc) and the presence of backbone-to-backbone hydrogen bonds (hbond_sr_bb and hbond_lr_bb). The energy terms are normalized before inputting into the deep learning neural network.

#### 2.2.4 Multi-distance and Orientations

To effectively describe the information of the protein structure, we use (i) inter-residue *C*_*β*_ distance map, (ii) *C*_*α*_ to tip atom distance map, (iii) tip atom to *C*_*α*_ distance map, (iv) tip atom to tip atom distance map, (v) sequence separation map, and (vi) inter-residue orientations (*φ,θ,ω*) defined by trRosetta (Yang et al., 2020). The sequence separation can be calculated as follows:

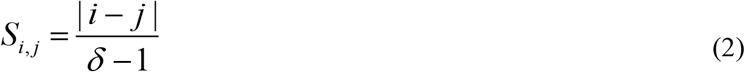

where *i* and *j* represent the residue index in the protein sequence and *δ*represents the normalizer, which is set to 100. The definition of inter-residue orientations can be found in Figure S1 in SI, and the tip atoms of each residue are listed in Table S3 in SI.

### 2.3 Network architecture

Figure 1(c) shows an overview of the network architecture. First, the 3D voxelization feature passes through a series of 3D convolution layers. The output tensor of 3D convolution layers is flattened into a 1D vector and concatenated with 1D features then goes through a 1D convolution layer. The output tensor of 1D convolution layer is transformed into a 2D tensor by horizontal and vertical striping then combined with other 2D tensor form feature maps. Second, the feature maps are input to a 2D convolutional layer and apply instance normalization. ELU activation is then used. The output is upsampled to 128 channels for the following ResNet operations in this work.

The ResNet operations contains trunk and arm residual blocks. The trunk part consists of 15 residual blocks, each of which is composed of three ELU activation layers, three 2D convolutional layers (dilation is applied with a cycling dilation size of 1,2,4,8 and 16), and three instance normalization layers. The network then branches to two arms of four residual blocks. The output layer distribution of the two arms consists of sigmod and softmax functions to predict *C*_*β*_ distance deviations for all residue pairs (referred to as deviation) and contact map with thresholded at 15Å. As shown in Figure S2 in SI, in each residual block, a “skipconnection” is made between the input and the output layers by skipping the intermediate layer. This connection works as an identity mapping, and its outputs are added to the output of all the layers that were previously stacked and passed to the activation phase of the last layer of the residual block.

### 2.4 Model training and lDDT calculation

lDDT is a superposition-free score that evaluates the local distance differences of all atoms in a protein structure, and it is well suited for assessing local protein model quality (Mariani *et al*., 2013). The deviation and contact map thresholded at 15Å predicted by our deep residual network can be used to calculate the lDDT score of each residue in a protein model. The lDDT score of the ith residue and the global lDDT score are defined as follows:

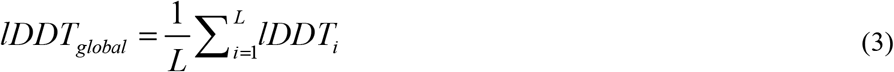

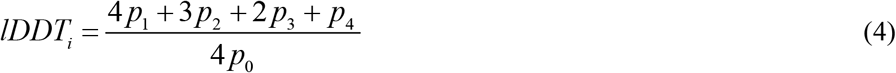

where *p*_0_ is the probability that the distance between the ith residue and other residues is within 15Å, and *p*_1_ is the probability that the absolute value of *C*_*β*_ distance deviation of all the residue pairs whose distance from the ith residue within 15Å is less than 0.5Å. In the same manner, *p*_2_, *p*_3_, and *p*_4_ are the probability that the absolute value of *C*_*β*_ distance deviation in all the residue pairs whose distance from the *i*th residue is within 15Å is 0.5Å−1.0Å, 1.0Å−2.0Å, and 2.0Å−4.0Å, respectively.

The weights in the network are initialized through Xavier initialization (Glorot et al., 2010) with weights drawn from the uniform distribution, and then obtained with the Adam method which is widely used by the neural network model (Kingma *et al*., 2014; Zhou *et al*., 2020; Jing *et al*., 2021; Li *et al*., 2021) at an initial learning rate of 0.0005. The multivariate cross-entropy loss function is used to evaluate the residual pair distance deviation loss, and the binary crossentropy loss function is adopted to evaluate the loss of the contact map with distance cutoff of 15Å. The mean square loss function is used to evaluate the lDDT loss of each residue. Then, the combination of distance error loss, mask loss, and lDDT loss is minimized as:

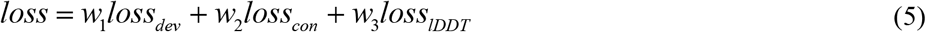

Where *loss*_*dev*_ is the distance deviation loss, *loss*_*dev*_ is the contact (thresholded at 15Å) loss, and *loss*_*lDDT*_ is the lDDT loss of each residue. The weights of each loss *w*_1_, *w*_2_ and *w*_3_ are 1, 1, and 10, respectively.

## 3 Results and discussions

Three evaluation indicators are used to evaluate the performance of the model quality assessment: (i) Correlation between the predictied and real score of protein model quality. Pearson, Spearman and Kendall’s Tau correlation coefficients are used to measure the average correlation between the predicted lDDT of each target model and the real lDDT, and the overall correlation between the predicted lDDT and the real lDDT of all target models (Bolboaca *et al*., 2006). The higher the correlation is, the better the performance is. (ii) Capability to find the best model. It is measured by using the lDDT difference (referred to as “average loss”) between the best model selected based on the predicted lDDT and the actual best model of the target protein. The lower the average loss, the better the performance. (iii) Capability to distinguish between good and poor models, which is measured by the area under the receiver operating characteristic (ROC) curve with an lDDT cutoff value of 0.6. The larger the area under the curve (AUC) of ROC, the better the capability is to distinguish between good and poor models (Ling *et al*., 2003).

### 3.1 Results of CASP targets

The performance of DeepUMQA is tested on 13,970 structure models composed of 51 CASP13 and 44 CASP14 targets and compared with the performance of state-of-the-art single-model quality assessment methods, namely, ProQ2 (Ray *et al*., 2012), ProQ3 (Uziela *et al*., 2016), ProQ3D (Uziela *et al*., 2017), VoroMQA (Olechnovic *et al*., 2017), Ornate (Pagès *et al*., 2019), and DeepAccNet (Hiranuma *et al*., 2021). The results of DeepAccNet are obtained by running the official source code locally, and the results of the other methods are obtained from the data archive on the official CASP website (https://predictioncenter.org/download_area/).

The performance of DeepUMQA and the compared methods on the CASP13 and CASP14 datasets are shown in Table 1. And the detailed results for each target are listed in Tables S8 and S9 in SI. In CASP13 and CASP14 databases, DeepUMQA consistently outperforms all of the compared methods in terms of all indicators. DeepUMQA has the highest global Pearson, Spearman, Kendall’s Tau correlation (0.8332, 0.8131, and 0.6231, respectively) and the average Pearson, Spearman, Kendall’s Tau correlation per target (0.7611, 0.7529, and 0.5743, respectively) in CASP13 test dataset. These values are much better than those of the second-best DeepAccNet (the 6 indicators have values of 0.8157, 0.7997, 0.6118, 0.7313, 0.7384, and 0.5635, respectively) in CASP13 dataset. In addition, the average lDDT loss of DeepUMQA is 0.0292, which is better than the value of all the compared methods. For the CASP14 dataset, the global Pearson, Spearman, Kendall’s Tau correlation of DeepUMQA are 0.7891, 0.7818, and 0.5854, respectively, which are also better than that of all the compared methods. The average Pearson, Spearman, Kendall’s Tau correlation of DeepUMQA are 0.7273, 0.7126, and 0.5324, respectively, which are also better than that of all the compared methods. In addition, DeepUMQA has the lowest average lDDT loss of 0.0473. To analyze the capability of DeepUMQA for distinguishing good models from poor models, we use all models in CASP13 and CASP14 datasets to perform a ROC analysis. Figure 4 shows the ROC curves with AUC. In the CASP13 and CASP14 datasets, the AUC value of DeepUMQA are 0.93804 and 0.90163, respectively, which are significantly higher than the value for the compared methods (ProQ2, ProQ3, ProQ3D, VoroMQA, Ornate, and DeepAccNet, respectively).. With regard to the correlation between the predicted and real scores of protein model quality, the capability to find the best model, and the capability to distinguish between good and poor models (three indicators), DeepUMQA is consistently superior to the compared state-of-the-art methods. We also find that among the compared methods, the deep learning-based approaches, such as ProQ3D, Ornate, and DeepAccNet, perform better than the SVM-based approaches, such as ProQ2 and ProQ3. This shows that protein model quality assessment is transitioning from traditional machine learning to deep learning-based methods.

**Table 1.** Performance of DeepUMQA and the compared single-model quality assessment methods on CASP13 and CASP14 datasets.

| Dataset | Method | Av.p | Av.s | Av.k | Gl.p | Gl.s | Gl.k | Av.loss |
| --- | --- | --- | --- | --- | --- | --- | --- | --- |
| CASP13 | **DeepUMQA** | **0.7611** | **0.7529** | **0.5743** | **0.8332** | **0.8131** | **0.6231** | **0.0292** |
|  | DeepAccNet | 0.7313 | 0.7384 | 0.5635 | 0.8157 | 0.7997 | 0.6118 | 0.0312 |
|  | ProQ3D | 0.7171 | 0.6883 | 0.5392 | 0.7825 | 0.7946 | 0.6007 | 0.0518 |
|  | ProQ3 | 0.6928 | 0.6745 | 0.5029 | 0.7717 | 0.7702 | 0.5838 | 0.0438 |
|  | ProQ2 | 0.6832 | 0.6512 | 0.4862 | 0.7687 | 0.7714 | 0.5829 | 0.0679 |
|  | VoroMQA | 0.7152 | 0.6824 | 0.5185 | 0.6723 | 0.6757 | 0.5048 | 0.0356 |
| CASP14 | **DeepUMQA** | **0.7273** | **0.7126** | **0.5324** | **0.7891** | **0.7818** | **0.5854** | **0.0473** |
|  | DeepAccNet | 0.7024 | 0.6936 | 0.5168 | 0.7778 | 0.7751 | 0.5814 | 0.0521 |
|  | Ornate | 0.7059 | 0.6524 | 0.4820 | 0.6869 | 0.6824 | 0.4978 | 0.0591 |
|  | ProQ3D | 0.6424 | 0.6068 | 0.4452 | 0.6743 | 0.6719 | 0.4838 | 0.0904 |
|  | ProQ2 | 0.6446 | 0.5919 | 0.4290 | 0.5401 | 0.5321 | 0.3835 | 0.0707 |
|  | ProQ3 | 0.6052 | 0.5177 | 0.3749 | 0.4610 | 0.4543 | 0.3341 | 0.0871 |

Note: Values in bold represent the best performance.
Av.p, Av.s, Av.k: Per-target average Pearson, Spearman and Kendall's Tau correlation with respect to real IDDT score.
Gl.p, Gl.s, Gl.k: Global Pearson, Spearman and Kendall's Tau correlation with respect to real IDDT score.
Av.loss: Per-target average loss with respect to real IDDT score. (The difference between the real IDDT scores of the best model selected from the predicted scores and the actual best model.)

**Fig. 4.**
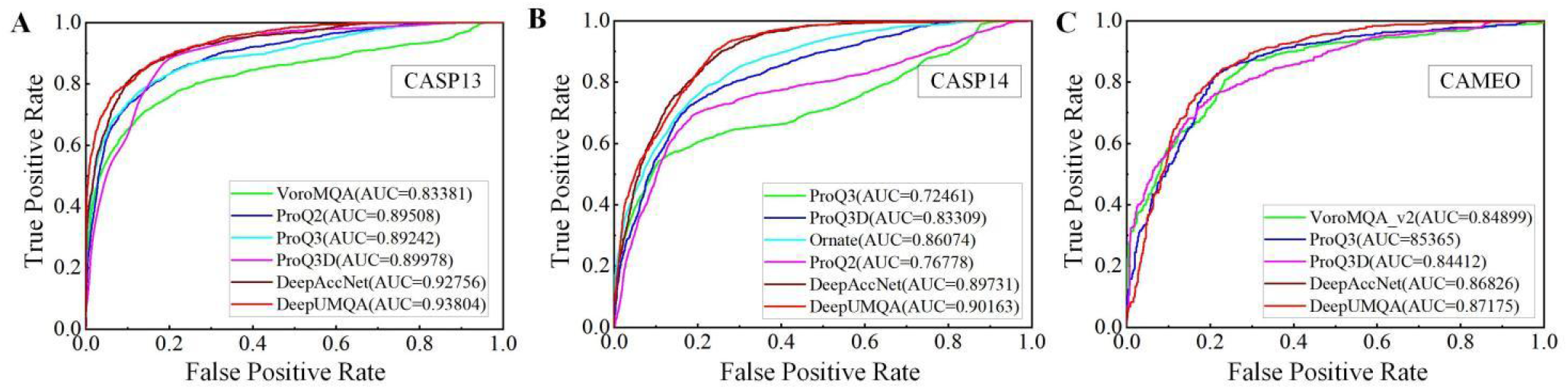
Receiver operating characteristic (ROC) curve of DeepUMQA and the compared single-model quality assessment methods on CASP13, CASP14 and CAMEO datasets. An lDDT cutoff value of 0.6 is used to distinguish good and poor models in (A) CASP13, (B) CASP14, and (C) CAMEO datasets respectively.

### 3.2 Results on CAMEO targets

The perfomance of DeepUMQA is also tested on 3,087 models for 195 CAMEO targets and compared with the performance of VoroMQA, ProQ3, ProQ3D, and DeepAccNet. In addition to DeepAccNet run locally with parameters set according to its paper, we directly obtain the quality assessment predictions submitted by the test methods from the data archive on the CAMEO official website (https://www.cameo3d.org/quality-estimation/).

Table 2 presents the performance of DeepUMQA and VoroMQA, ProQ3, ProQ3D, and DeepAccNet on the CAMEO dataset. Given that the CAMEO dataset has only a small number of models per target (about 16 models/target), the average correlation and loss of each target have no obvious statistical significance. Hence, we only use three global correlations as evaluation indicators. The experimental results show that DeepUMQA achieves the highest global Pearson (0.7522), Spearman (0.7717) and Kendall’s Tau correlation (0.5822) compared with the other four state-of-the-art methods, which are 3.00%, 1.97%, and 2.90% better than the best results of the compared methods, respectively. We also perform ROC analysis on all models in the CAMEO dataset to analyze the capability of the various single-model quality assessment methods to distinguish good models from poor ones. Figure 4 (C) shows the ROC curves with AUC values. We find that DeepUMQA is still optimal, and its AUC is 0.87175, which is 0.40% better than the best methods of the compared methods (DeepAccNet with AUC of 0.86826). In the same manner, the deep learning-based approaches perform better than the SVM-based approaches.

**Table 2.** Performance of DeepUMQA and the compared single-model quality assessment methods on the CAMEO dataset.

| Dataset | Method | Gl.p | Gl.s | Gl.k |
| --- | --- | --- | --- | --- |
| CAMEO | **DeepUMQA** | **0.7522** | **0.7717** | **0.5822** |
|  | DeepAccNet | 0.7303 | 0.7568 | 0.5658 |
|  | VoroMQA | 0.5185 | 0.7096 | 0.5177 |
|  | ProQ3 | 0.4459 | 0.6729 | 0.4857 |
|  | ProQ3D | 0.6467 | 0.7143 | 0.5269 |

Note: Values in bold represent the best performance.
Gl.p, Gl.s, Gl.k: Global Pearson, Spearman and Kendall's Tau correlation with respect to the real IDDT score.

### 3.3 Effect of USR

To verify the effect of the USR feature, we train three comparative versions by using the same dataset and parameters as DeepUMQA, the three versions are DeepUMQA^1^ (DeepUMQA without voxelization and USR features), DeepUMQA^2^ (DeepUMQA without the voxelization feature), and DeepUMQA^3^ (DeepUMQA without the USR feature).

The performance of different versions of DeepUMQA on the CASP13, CASP14, and CAMEO test datasets is shown in Table 3, and the ROC curve analysis is shown in Figure 5. The experimental results show that compared with DeepUMQA^1^ without voxelization and USR features, DeepUMQA^2^ with the USR feature has higher Pearson, Spearman, and Kendall’s Tau correlations and lower average lDDT loss. The AUC of DeepUMQA^2^ is also better than that of DeepUMQA^1^. Similarly, the overall performance of DeepUMQA^3^ is improved by the use of voxel features compared with DeepUMQA^1^. Both USR and voxelization features can improve the performance of model quality assessment to some extent,and the performance of DeepUMQA^2^ and DeepUMQA^3^ is comparable on all the test datasets. This result shows that they can both reflect the structural information of residues. Notably, the use of both voxelization and USR features (i.e., DeepUMQA) improves the performance of model quality assessment, and almost all performance indicators are better than those for the three other versions on the three test datasets. This funding further indicates that USR and voxelization features can complement each other and reflect the topological and local structure information of residues comprehensively. In summary, all of the results emphasize the advantages of using the USR feature for model quality assessment, that is, model quality assessment predictors with the USR feature have a stronger correlation between the predicted and real score of models, and better capability to find the best models and distinguish good models from poor models.

**Table 3.** Performance comparison of different versions of DeepUMQA on CASP13, CASP14, and CAMEO datasets.

|  |  | DeepUMQA<sup>1</sup> | DeepUMQA<sup>2</sup> | DeepUMQA<sup>3</sup> | **DeepUMQA** |
| --- | --- | --- | --- | --- | --- |
| CASP13 | Av.p | 0.6955 | 0.7407 | 0.7313 | **0.7611** |
|  | Av.s | 0.7267 | 0.7407 | 0.7384 | **0.7529** |
|  | Av.k | 0.5547 | 0.5836 | 0.5635 | **0.5743** |
|  | Gl.p | 0.7908 | 0.8046 | 0.8157 | **0.8332** |
|  | Gl.s | 0.7853 | 0.7940 | 0.7997 | **0.8131** |
|  | Gl.k | 0.5939 | 0.6032 | 0.6118 | **0.6231** |
|  | Av.loss | 0.0414 | 0.0226 | 0.0312 | **0.0292** |
| CASP14 | Av.p | 0.6750 | 0.7025 | 0.7024 | **0.7273** |
|  | Av.s | 0.6766 | 0.6966 | 0.6936 | **0.7126** |
|  | Av.k | 0.4973 | 0.5170 | 0.5168 | **0.5324** |
|  | Gl.p | 0.7312 | 0.7563 | 0.7778 | **0.7891** |
|  | Gl.s | 0.7276 | 0.7557 | 0.7751 | **0.7818** |
|  | Gl.k | 0.5351 | 0.5621 | 0.5814 | **0.5854** |
|  | Av.loss | 0.0524 | 0.0494 | 0.0521 | **0.0473** |
| CAMEO | Gl.p | 0.6383 | 0.6664 | 0.7303 | **0.7522** |
|  | Gl.s | 0.6757 | 0.7091 | 0.7568 | **0.7717** |
|  | Gl.k | 0.4898 | 0.5221 | 0.5658 | **0.5822** |

Note: Values in bold represent the performance of DeepUMQA.
DeepUMQA<sup>1</sup>: DeepUMQA without both USR and voxelization features.
DeepUMQA<sup>2</sup>: DeepUMQA without the voxelization feature.
DeepUMQA<sup>3</sup>: DeepUMQA without the USR feature.

**Fig. 5.**
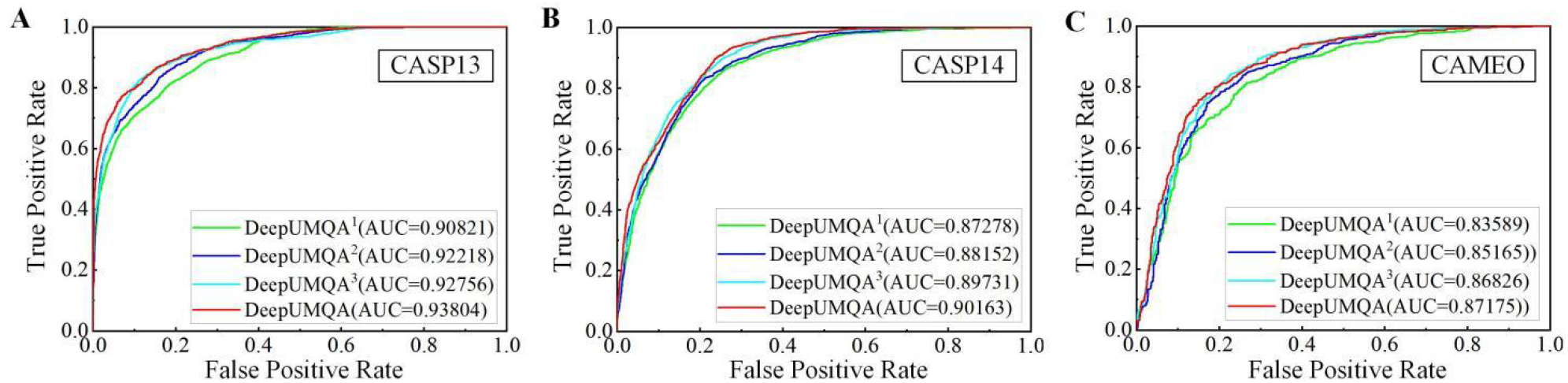
Receiver operating characteristic (ROC) curve of DeepUMQA and its different versions (DeepUMQA^1^, DeepUMQA^2^, and DeepUMQA^3^) on CASP13, CASP14, and CAMEO datasets. An lDDT cutoff value of 0.6 is used to distinguish good and poor models in (A) CASP13, (B) CASP14, and (C) CAMEO datasets respectively. DeepUMQA^1^, DeepUMQA^2^, and DeepUMQA^3^ represent DeepUMQA without USR and voxelization features, DeepUMQA without the voxelization feature, and DeepUMQA without the USR feature, respectively.

### 3.4 Qualitative Analysis

Figure 6 shows a quantitative analysis of the quality evaluation of target T1046s2 by using DeepUMQA. Figure 6(B) shows a comparison of the predicted lDDT and real lDDT for all test models of target T1046s2. They have a strong correlation, especially when the model has high accuracy with lDDT>0.5. The predicted Pearson correlation with the real lDDT is 0.8271, the Spearman correlation is 0.8204, and Kendall’s tau correlation is 0.6405. Furthermore, the best model selected based on our predicted lDDT score is consistent with the best model in the actual model pool (the sky-blue dot). Figure 6(C) shows the ROC curve of all test models of target T1046s2, where AUC is 0.98366, indicating the capability to distinguish good and poor models is excellent. Figure 6(D) shows the model assessment results of three randomly selected models with different qualities. The first row shows the structure of the three models and their corresponding predicted and real global model qualities, and the second row shows the comparison of the real lDDT (shown in gray) and predicted lDDT (shown in different colors) in each residue. Although differences exist between the predicted lDDT and the real lDDT in several residues, we observe that DeepUMQA can correctly capture the change trend of the lDDT of residue under models with different accuracies, which is important for correcting the mis-modeled regions of the model and further model refinement.

**Fig. 6.**
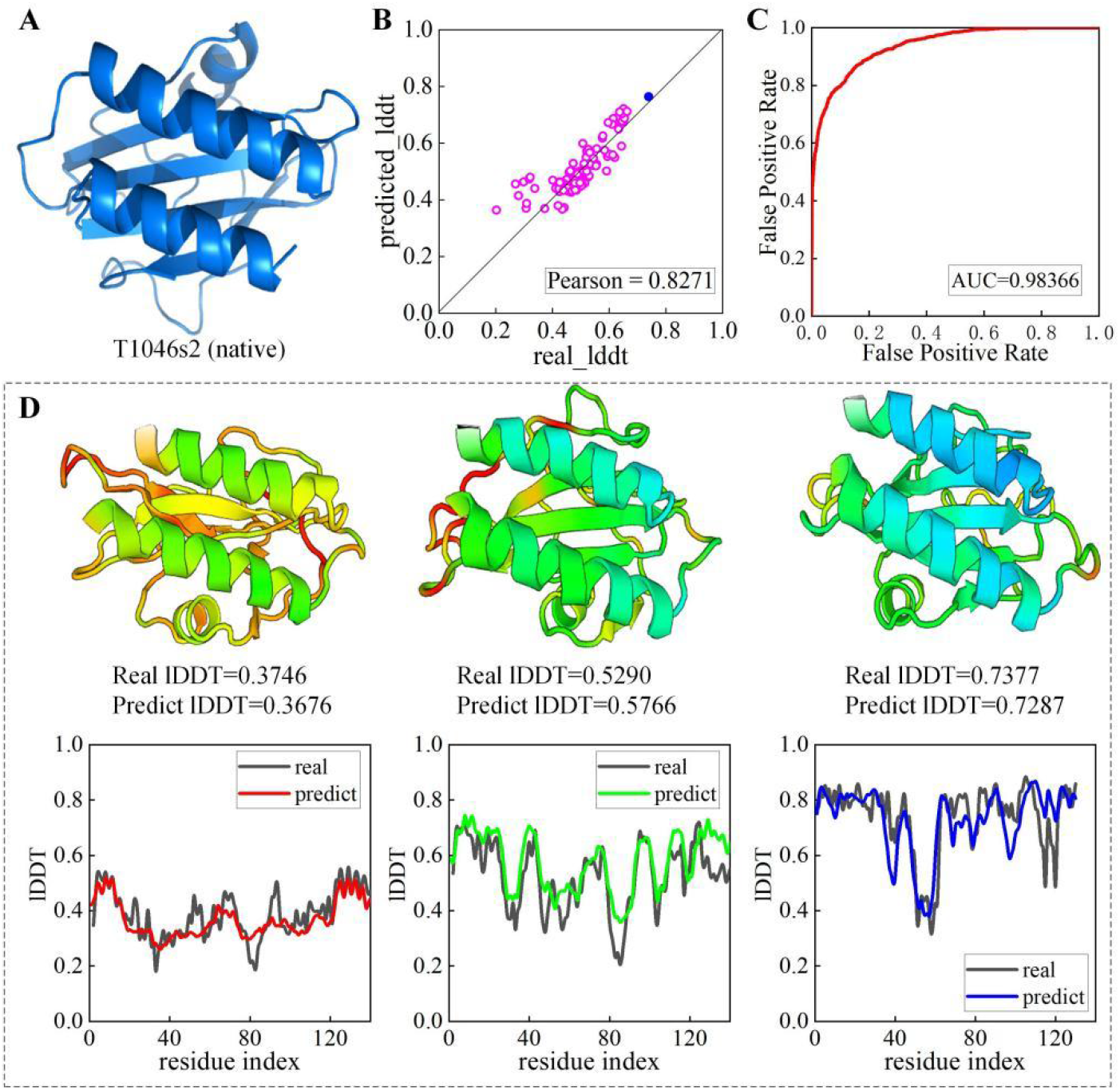
Qualitative analysis of the performance of DeepUMQA on CASP14 target T1046s2. (A) The experimental structure of the target T1046s2. (B) Comparison of the predicted lDDT and real lDDT for all test models of target T1046s2 (sky-blue solid point represents the best model). (C) The ROC curve of all test models of target T1046s2. (D) Results of model assessment of three randomly selected models of target T1046s2 with different quality.

## 4 Conclusion

We developed a USR-based residue-level single-model quality assessment method called DeepUMQA. Residue-level USR and voxelization features, amino acid sequence properties and secondary structure, Rosetta energy terms, and distances and orientations extracted from the structure model were used to describe the model information. These features were fed into the deep residual neural network to predict the quality of the protein structure model. The performance of DeepUMQA was extensively tested on 51 CASP13, 44 CASP14, and 195 CAMEO targets, which included 17,057 model structures. The experimental results showed that the performance of DeepUMQA was better than that of the state-of-theart single-model quality assessment methods, including ProQ2, ProQ3, ProQ3D, Ornate, VoroMQA, and DeepAccNet. The USR feature describes the residue-level topological structure information, and it is complementary to the voxelization feature that describes the local structure information of the residues. As a result, the residue structure information is comprehensively characterized, and the accuracy of model assessment is considerably improved. With the rapid development of protein structure prediction, the dynamic combination of model quality assessment and folding process could be a future research direction.

